# Risk stratification of allogeneic stem cell recipients with respect to the potential for development of GVHD via their pre-transplant plasma lipid and metabolic signature

**DOI:** 10.1101/475244

**Authors:** Daniel Contaifer, Catherine H Roberts, Naren Gajenthra Kumar, Ramesh Natarajan, Bernard J Fisher, Kevin Leslie, Jason Reed, Amir A Toor, Dayanjan S Wijesinghe

## Abstract

The clinical outcome of allogeneic hematopoietic stem cell transplantation (SCT) is strongly influenced from the complications arising during the post-transplant immune restoration and has been well studied and described. However, the metabolic status of the recipient pre-transplant also has the potential to influence this outcome and has never been studied before and has the potential to enable risk stratification with respect to the development of transplant associated complications such as graft vs. host disease (GVHD). In order to better understand this aspect of transplant related complications we investigated the pre-transplantation metabolic signature to assess the possibility of pre-transplant risk stratification. This pilot study was composed of 14 patients undergoing myeloablative conditioning followed by either HLA matched related, unrelated donor, or autologous stem cell transplantation. Blood samples were taken prior to transplant and the plasma was comprehensively characterized with respect to its lipidome and metabolome via LCMS and GCMS. The results indicated a significantly pro-inflammatory metabolic profile in patients who eventually developed Graft vs. Host Disease (GVHD). The data revealed 5 potential pre-transplant biomarkers (1-monopalmitin, diacylglycerol (DG) 38:5, DG 38:6, 2-aminobutyric acid, and fatty acid (FA) 20:1) that demonstrated high sensitivity and specificity towards predicting post-transplant GVHD development. The predictive model developed demonstrated an estimated predictive accuracy of risk stratification of 100%, with an Area under the Curve of the ROC of 0.995 with 100%. The likelihood ratio of 1-monopalmitin (infinity), DG 38:5 (6.0) and DG 38:6 (6.0) also demonstrated that a patient with a positive test result for these biomarkers pre-transplant will likely have very high odds of developing GVHD post-transplant. Collectively the data demonstrates the possibility of using pre-transplant metabolic signature for risk stratification of SCT recipients with respect to development of GVHD.

## Introduction

Transplantation of hematopoietic progenitors from an HLA matched donor is a curative procedure for many patients with hematologic malignancies and disorder of hematopoiesis. Graft vs. host disease (GVHD) is a frequently observed complication of stem cell transplantation (SCT) which contributes to transplant related mortality and adversely impacts clinical outcomes following transplantation. GVHD after allogeneic HSCT is a reaction of donor immune cells to recipient tissues. An inflammatory cascade triggered by the preparative regimen causes activated donor T cells to damage epithelial cells. About 35%–50% of HSCT recipients will develop acute GVHD. It is mediated by donor-derived T cells responding to minor histocompatibility antigens encountered in the recipient. The T cell encounter alloantigens, undergo activation, perform functions such as cytokine secretion (IL-2, IL-4, IL-10, IL-12, IL-17 and interferon gamma by helper T cell subsets) and target lysis (granzyme and perforin secretion by cytotoxic T cells). These functions are accompanied by significant metabolic adaptations in the T cells, including increased glycolysis and oxygen consumption as well as cytokine production mentioned earlier^1^. As an example, higher levels of GLUT 1 expression have been observed in activated T cells, suggesting increased metabolic & biosynthetic rates^2^. Supporting this, a correlation has been shown between intracellular ATP concentration in T cells and severity of clinical GVHD in humans and between increasing glycolysis and GVHD in murine models^3,4^. These results suggest that T cell activation and consequent metabolic and biosynthetic, and logically, mass changes may correlate with significant functional events in the immune reconstitution process following SCT, as donor-derived T cells experience a new antigenic milieu post-transplant.

Just like metabolic changes in the T cell are crucial to the onset of immune reactions, the metabolic milieu in which the T cells find themselves, influences their function. In this respect, the lipid molecules constitute a family of immensely important functional mediators present in the metabolic milieu. The effects of some lipids on the T cells have been studied recently. These effects include lysophosphatidylserine (lysoPS) mediated suppression of IL-2 production and suppression of T cell proliferation^5^. This effect is mediated via LPS3/G protein coupled receptor 174 which triggers IL-2 production in CD4+ T cell. Another enzyme, acid sphingomyelinase (ASMase) generates ceramide, and modulates signaling cascades involving CD3 and CD28. It is involved in Th1 and Th17 responses through its effect on STAT 3 and mTOR^6^. An acid sphingomyelinase deficient mouse model experiences attenuation of GVHD^7^. Along the same lines, leukotriene C4 has been shown to be important for airway inflammation when administered to murine models along with IL33^8^. Consistent with such results are the observations that T cells in acetyl-CoA carboxylase deficient mice are resistant to induction of GVHD^9^. T effector cells have been shown to increase their reliance on fatty acid metabolism during GVHD as well ^10,11^, for example, Prostaglandin E2 (PGE_2_) has been implicated in modulating T cell effects of mesenchymal stromal cells on T cell populations^12^ and PGE_2_ priming of T cells reduces GVHD when localized to the site of alloreactivity^13^. Further, bone marrow stromal cells also exert an ameliorating influence on GVHD through indoleamine 2,3-dioxygenase (IDO) and PGE_2_ expression^14^. These lipid mediated effects have been targeted in the treatment of GVHD, for instance, the effectiveness of leukotriene inhibitor monteleukast has long been recognized in managing GVHD of the lung^15,16^. Additionally, prostaglandins have been studied in GVHD prevention strategies^17^, particularly PGE_2_ ^13^. A leukotriene inhibitor, eicosapentanoic acid (EPA) has also been studied in the treatment of GVHD as well as prophylaxis^18,19^. Prostaglandin mediators FT1050 (16,16-dimethyl PGE2) and FT4145 (dexamethasone) are also being studied in clinical trials of GVHD prophylaxis using ex-vivo modification of the allograft^20,21^. These observations make it crucial to gain an understanding regarding the lipid and metabolic changes that come about following dose intensive, myeloablative conditioning, and how the metabolome and the lipidome might impact T cell function following SCT. Modern methods of lipidomics and metabolomics allows us to study such changes in detail^22–26^.

Recent studies have demonstrated that a dynamical systems model of T cell reconstitution may allow understanding alloreactivity in terms of antigenic differences between donors and recipients and ensuing T cell responses^27–29^. This model incorporates the idea that when T cells become activated and undergo metabolic changes upon antigen encounter, they may increase mass, and this appears to be the case when T cell masses are measured following allogeneic SCT^30^. This change in mass is likely driven by the cytokine and chemokine milieu in these patients, which is likely to be altered because of conditioning chemotherapy and radiation induced tissue damage. Further chemokine release is mediated by the effects of inflammation due to infections and endothelial injury. These changes in the post-transplant milieu may be investigated through the study of the metabolomic and lipidomic profile. In this paper we describe the lipidomic and metabolomic profiles of patients undergoing myeloablative conditioning and stem cell transplantation obtained prior to the infusion of stem cell graft and potential towards risk stratification with respect to the development of the GVHD.

## Materials and Methods

### Patients

Patients were enrolled prospectively in a Virginia Commonwealth University (VCU) Institutional Review Board (IRB)-approved observational study; patients provided written informed consent prior to enrollment. Patients underwent myeloablative conditioning followed by either HLA matched related (MRD), unrelated donor (URD), or autologous stem cell transplantation (auto). Conditioning regimens utilized included busulfan/cyclophosphamide (5), busulfan or melphalan with fludarabine (3 & 2 respectively), carmustine, etoposide, cytarabine and melphalan (3), total body irradiation and cyclophosphamide, melphalan (1 each). HLA matching was at the allelic level; allogeneic HCT recipients received ATG, calcineurin inhibitors and cellcept or methotrexate for GVHD prophylaxis. Blood samples were drawn prior to transplant, and on days 0, 14, 30, and 60 post-SCT. Blood was processed for plasma collection and frozen at −80°C until mass spectroscopy based metabolomic and lipidomic analysis. Given the small patient cohort, acute and chronic GVHD data were pooled, Glucksberg and NIH Consensus criteria were used to diagnose and stage GVHD.

### Lipid and metabolite extraction for LC-MS/MS analyses

Blood plasma lipids extraction was carried out using a biphasic solvent system of cold methanol, methyl tertiary butyl ether (MTBE), and water with some modifications (Matyash et al. 2008). In detail, 225 µL of cold methanol containing a mixture of odd chain and deuterated lipid internal standards [lysoPE(17:1), lysoPC(17:0), PC(12:0/13:0), PE(17:0/17:0), PG(17:0/17:0), sphingosine (d17:1), d7 cholesterol, SM(17:0), C17 ceramide, d3 palmitic acid, MG(17:0/0:0/0:0), DG(18:1/2:0/0:0), DG(12:0/12:0/0:0), and d5 TG(17:0/17:1/17:0)] was added to a 20 µL blood plasma aliquot in a 1.5 mL polypropylene tube, and then vortexed. Next, 750 µL of cold MTBE was added, followed by vortexing and shaking with an orbital mixer. Phase separation was induced by adding 188 µL of MS-grade water. Upon vortexing (20 s) the sample was centrifuged at 12,300 rpm for 2 min. The upper organic phase was collected in two 300 µL aliquots and evaporated with a vapor trap. Dried extracts were resuspended using 110 µL of a methanol/toluene (9:1, v/v) mixture containing CUDA (50 ng/ml; internal standard for quality control of injection) with support of vortexing (10 s), and centrifuged at 800 rpm for 5 min, followed by transferring 100 uL of the supernatant into autosampler vial with an insert. The lower polar layer was collected and an aliquot of 125 µL was evaporated to dryness in a SpeedVac and resuspended in acetonitrile for polar metabolite analysis via HILIC LC-MS/MS method.

### Metabolomics:GC-MS metabolite extraction

30μl of plasma sample was added to a 1.0 mL of pre-chilled (−20°C) extraction solution composed of acetonitrile, isopropanol and water (3:3:2, v/v/v). Sample were vortexed and shaken for 5 min at 4°C using the Orbital Mixing Chilling/Heating Plate. Next, the mixture was centrifuged for 2min at 14000 rcf. 450μL of the supernatant was dried with cold trap concentrator. The dried aliquot was then reconstituted with 450 μL acetonitrile:water (50:50) solution and centrifuged for 2 min at 14000 rcf. The supernatant was transferred to a polypropylene tube and subjected to drying in a cold trap. The process of derivatization began with addition of 10 μL of 40 mg/mL Methoxyamine hydrochloride solution to each dried sample and standard. Samples were shaken at maximum speed at 30 ^o^C for 1.5 hours. Then, 91 μL of MSTFA + FAME mixture was added to each sample and standard and capped immediately. After shaking at maximum speed at 37°C, the content was transferred to glass vials with micro-serts inserted and capped immediately.

### Lipids: LC-MC/MC conditions

Untargeted lipid analysis was undertaken with Sciex TripleTOF 6600 coupled to Agilent 1290 LC. Lipids were separated on an Acquity UPLC CSH C18 column (100 × 2.1 mm; 1.7 µm) (Waters, Milford, MA, USA). The column was maintained at 65 °C and the flow-rate of 0.6 mL/min. The mobile phases consisted of (A) 60:40 (v/v) acetonitrile:water with 10 mM ammonium acetate and (B) 90:10 (v/v) isopropanol:acetonitrile with 10 mM ammonium acetate. The separation was conducted following a stepwise gradient: 0-2 min 15-30% (B), 2-2.5 min 48% (B), 2.5-11 min 82% (B), 11-11.5 min 99% (B), 11.5-12 min 99% (B), 12-12.1 15% (B), 12-14 min 15% (B). Negative and positive electrospray ionization (ESI) modes were applied with nitrogen serving as the desolvation gas and the collision gas. The parameters for detection of lipids were as follows: Curtain Gas: 35; CAD: High; Ion Spray Voltage: 4500 V; Source Temperature: 350°C; Gas 1: 60; Gas 2: 60; Declustering Potential: +/-80V, and collision energies +/-10.

### Metabolites HILIC: LC-MS/MS conditions

Detection of water soluble plasma metabolites was carried out on Sciex TripleTOF 6600 coupled to Agilent 1290 LC. Analytes were separated on an Acquity UPLC BEH Amide Column, 130Å, 1.7 µm, 2.1 mm X 150 mm (Waters, Milford, MA, USA). The column was maintained at 45°C and the flow-rate of 0.4 mL/min. The mobile phases consisted of (A) water with 10 mM ammonium formate, 0.125% formic acid, and (B) acetonitrile:water 90:10 (v/v) with 10 mM ammonium formate, 0.125% formic acid. The analytes separation was conducted following a stepwise gradient: 0-2 min 100% (B), 2-7.7 min 70% (B), 7.7-9.5 min 40% (B), and 9.5-10.25 min 30% (B), and 410.25-12.75 min 100% (B), 12.75-16.75 100% (B). A sample volume of 1 µL and 3 µL were injected for positive and negative mode, respectively. Negative and positive electrospray ionization (ESI) modes were applied with nitrogen serving as the desolvation gas and the collision gas. The parameters for detection of lipids were as follows: Curtain Gas: 35; CAD: High; Ion Spray Voltage: 4500 V; Source Temperature: 300°C; Gas 1: 60; Gas 2: 60; Declustering Potential: +/-80V, and collision energies +/-10.

### Metabolites: GC-MS conditions

A Leco Pegasus IV time of flight mass spectrometer coupled with Agilent 6890 GC equipped with a Gerstel automatic liner exchange system (ALEX) that included a multipurpose sample (MPS2) dual rail, and a Gerstel CIS cold injection system (Gerstel, Muehlheim, Germany) was used to complement HILIC metabolite analysis. The transfer line was maintained at 280 °C. Chromatography separation was achieved on a 30 m long, 0.25 mm i.d. Rtx-5Sil MS column (0.25 μm 95% dimethyl 5% diphenyl polysiloxane film) with the addition of a 10 m integrated guard column was used (Restek, Bellefonte PA) with helium (99.999%; Airgas, Radnor, PA, U.S.A.) at a constant flow of 1 mL/min. The oven temperature was held constant at 50°C for 1 min and then ramped at 20°C/min to 330°C at which it is held constant for 5 min. The GC temperature program was set as follows: 50°C to 275°C final temperature at a rate of 12 °C/s and hold for 3 minutes. The injection volume was 1 μL in splitless mode at 250 °C. Electron impact ionization at 70V was employed with an ion source temperature of 250°C. The scan mass ranged from 85 to 500 Da with acquisition rate of 17 spectra/second.

### Statistical analysis

Prior to statistical analysis, metabolomic and lipidomic data were subjected to preprocessing as follows. Data was first normalized by a variant of a ‘vector normalization’ by calculating the sum of all peak heights for all identified metabolites for each sample and thereafter normalizing each compound by the total average of the sum. Multivariate statistical analysis tends to focus on metabolites with high intensities. To avoid this tendency, log scaling was applied to reduce the effect of large peaks and scale the data into a more normally distributed pattern. Pareto scaling, which uses the square root of the standard deviation as the scaling to change the emphasis from metabolites with high concentrations to those with moderate or small abundances, was also used for analyzing parameters with large variation.

Patient cohort was divided into patients that developed either acute or chronic GVHD and patients that did not develop GVHD. Since the analysis was performed before transplantation, autologous stem cell transplant patients that did not develop GVHD were grouped with allograft recipients that also did not develop GVHD. Partial Least Squares Discriminant Analysis (PLS-DA) was used to create a bilinear model to fit the data (Figure 1). Multivariate statistical methods as PLS-DA have been introduced to reduce the complexity of metabolic spectra and help identify meaningful patterns in high-resolution mass spectrometric data. In this method, the PLS-DA scores can be filtered through calculation of the variable importance in projection (VIP) and used to estimate the contribution of lipids and metabolites for class separation. The top 20 more important variables in the PLS-DA model was selected for further investigation (Figure 1).

**Figure 1.**
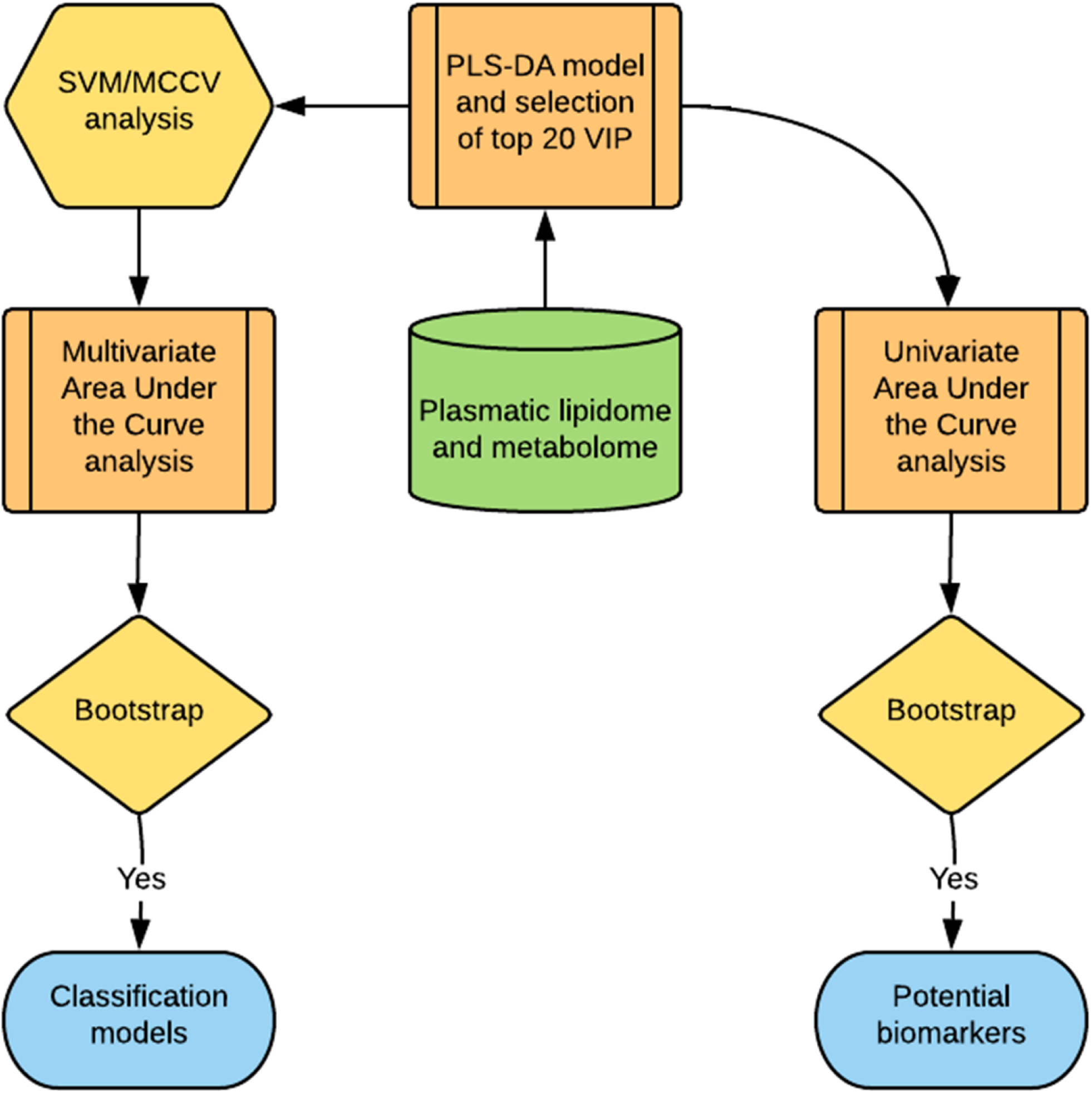
Statistical approach used for the identification of potential lipid and metabolite based biomarkers for the prediction of the onset of GVHD following BMT. A supervised statistical approach in the form of PLS-DA was applied to the consolidated metabolomic and lipidomic data as a first step to find potential biomarkers and to detect the presence of class separation, if any, between GVHD and no-GVHD patients. The top 20 more important variables that separate the groups was selected by Variable Importance in Projection (VIP). A Support Vector Machine (SVM) used these VIP variables to further find the binary classification of patients in the two groups. The result was cross-validated with Monte Carlo Cross-Validation (MCCV) and used in a multivariate AUCROC analysis to select the best model based on low dimensionality and high accuracy. Model estimates were validated with bootstrap CI. The best model was used as reference to select potential biomarkers of GVHD through univariate analysis of the top 20 VIP passing the criteria of high AUCROC estimate and low t-test p-value and validated with bootstrap CI.

Because of the relatively small sample size, cross validation was employed to further evaluate the classification model performance. An algorithm based on Support Vector Machine (SVM) was used to identify the top 20 variable importance in projection (VIP) to further select the best hyperplane that represents the largest separation between the two groups (Figure 1). This method was coupled with Monte-Carlo Cross Validation (MCCV) through balanced subsampling to support the validation in the small sample cohort (Figure 1). In each MCCV, two thirds of the samples are used to evaluate the feature importance and the left out one third is used as test population. The SVM and MCCV allows a multivariate Area under the Curve of the Receiver Operating Characteristic (AUCROC) analyses to estimate the success of the classification model (Figure 1), creating several AUCROC models to test performances with different numbers of predictors. This procedure was repeated multiple times to calculate the performance estimates and build a confidence interval for each model. Based on the performance in the Multivariate AUCROC the best number of predictors is a reference for selection of potential biomarkers^31^.

Univariate AUCROC analyses was used to find potential biomarkers with sufficient power to separate the groups (Figure 1). The result shows the t-test p-value, the AUCROC value with the confidence interval computed using 500 bootstrap replications. The criteria to choose the stronger potential biomarkers for GVHD was to select compounds with the highest AUCROC performance and lowest p-value (p<0.05). The calculated optimal cutoff was used to estimate the associated sensitivity and specificity values. Positive and negative likelihood ratios are calculated from the sensitivity and specificity output. The data was analyzed using MetaboAnalyst 3.5 maintained by Xia Lab at McGill University.

## Results

A total of 14 patients were accrued in this study (Table 1). Of these, 10 underwent a myeloablative allograft and 4 underwent an autologous SCT. Cohort was composed of 7 match related donors, 3 unrelated donors, and 4 autologous stem cell transplant recipients, with mean age of 50 (±10) years old, 57% women and 43% male, and 11 Caucasians and 3 African Americans.

**Table 1.** Demographic data for the cohort study. BMI = Body Mass Index; MRD = match related donor; URD = unrelated donor; auto = autologous; C = Caucasian; AA = African American; HTN = hypertension; MI = myocardial infarction; n/a = not applicable. Bu= Busulfan; Cy= Cytoxan; Cyls= cyclosporine; MTX= methotrexate; Tac= tacrolimus; MMF= mycopehnolate mofetil; TBI= total body irradiation; Flu= fludarabine; Mel= melphalan; BEAM= carmustine, etoposide, cytarabine and melphalan

| Donor type | Age | Gender | Race | BMI | Prep regimen; GVHD prophylaxis | GVHD (grade/severity) |
| --- | --- | --- | --- | --- | --- | --- |
| MRD | 60 | Female | C | 21.41 | Bu/Cy; Cyls/MTX | Acute (1) |
| URD | 50 | Female | C | 24.42 | Bu/Cy; Tac/MMF | Acute (4) |
| URD | 31 | Female | C | 34.99 | Bu/Cy; Tac/MTX | Chronic (mod) |
| URD | 40 | Female | C | 42.08 | TBI/Cy; Tac/MTX | Acute (2) |
| URD | 57 | Male | C | 22.09 | Bu/Flu; Tac/MMF | Acute (2) |
| URD | 50 | Male | C | 26.13 | Flu/Mel; Tac/MTX | Acute (1) |
| MRD | 59 | Male | C | 21.80 | Bu/Flu; Cyls/MMF | Chronic (mod) |
| URD | 50 | Female | C | 28.85 | Bu/Cy; Tac/MTX | no |
| MRD | 63 | Female | C | 27.81 | Bu/Flu; Cyls/MTX | no |
| URD | 66 | Male | C | 29.45 | Flu/Mel; Tac/MMF | no |
| auto | 48 | Female | C | 17.60 | BEAM; n/a | n/a |
| auto | 35 | Female | AA | 25.04 | Mel 200; n/a | n/a |
| auto | 44 | Male | AA | 45.22 | BEAM; n/a | n/a |
| auto | 40 | Male | AA | 51.22 | BEAM; n/a | n/a |

Following preprocessing and filtering to remove low quality data, in the individual patient data sets, the final aggregate, analyzable data set consisted of 225 plasma lipids and 139 non lipid small molecule metabolites derived from the patients. The statistical approach used for data analysis (Figure 1) was designed to provide meaningful and validated results optimized for the small sample size of the cohort, aiming to find potential biomarkers of GVHD to support future studies. To obtain an adequately powered sample, patients who did not experience GVHD following an allograft were combined with patients who underwent an autologous SCT in this pilot project.

### Pre-transplant plasma lipid and metabolite profiles reveals class separation between those patients who ultimately developed GVHD and those who did not

To estimate the potential of the lipids and metabolites to predict predisposition towards the development of GVHD in the SCT patients, pre-transplant plasma lipid and metabolite data were analyzed via PLS-DA. The degree of separation of patients with future GVHD against patients with no-GVHD was visualized by the scores plot of the two principal components. The distance of class separation suggests that metabolic variation may be used to predict GVHD in patients undergoing SCT (Figure 2).

**Figure 2.**
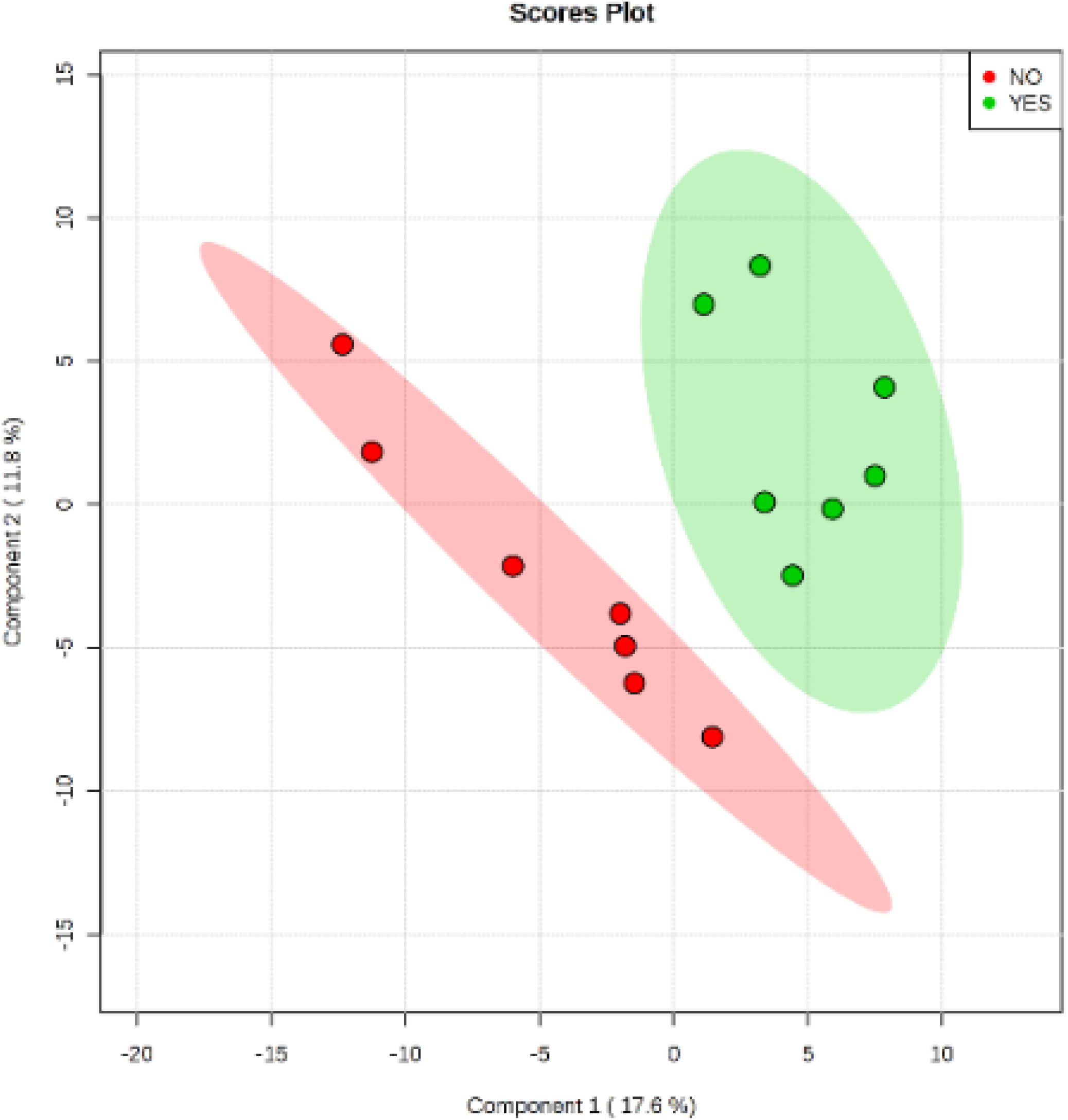
Supervised method PLS-DA found the direction of maximum covariance between the data set and the class membership, extracting features in form of latent variables. The model separation of GVHD (YES) and no-GVHD (NO) pre-transplant indicates metabolic differences for prediction of GVHD phenomena with no overlap in the 95% confidence intervals (shaded areas).

The top 20 variables that contributed to the separation observed in the PLS-DA model were represented by 2 metabolites and 18 lipids (Table 2). These characterized the metabolic variation present in the patients pre-transplant, and correlates strongly with GVHD development post-transplant. Compared to patients with no-GVHD, GVHD patients had decreased levels of 2-aminobutyric acid, hexose, monounsaturated fatty acids (14:1, 16:1, 18:1, 19:1 and 20:1) and poly-unsaturated fatty acid (20:3), as well as plasmenyl-ethanolamine (PE(p-34:1) or PE(o-34:2)). Further, GVHD patients presented elevation of monoacylglycerols (1-monoolein and 1-monopalmitin), diacylglycerols (38:5 and 38:6), along with elevated lysophosphocholine (14:0, 20:0), phosphocholines (28:0, 14:0/16:1, 16:0/18:3) and phosphoethanolamines (16:0/18:1, 18:0/22:5).

**Table 2.**
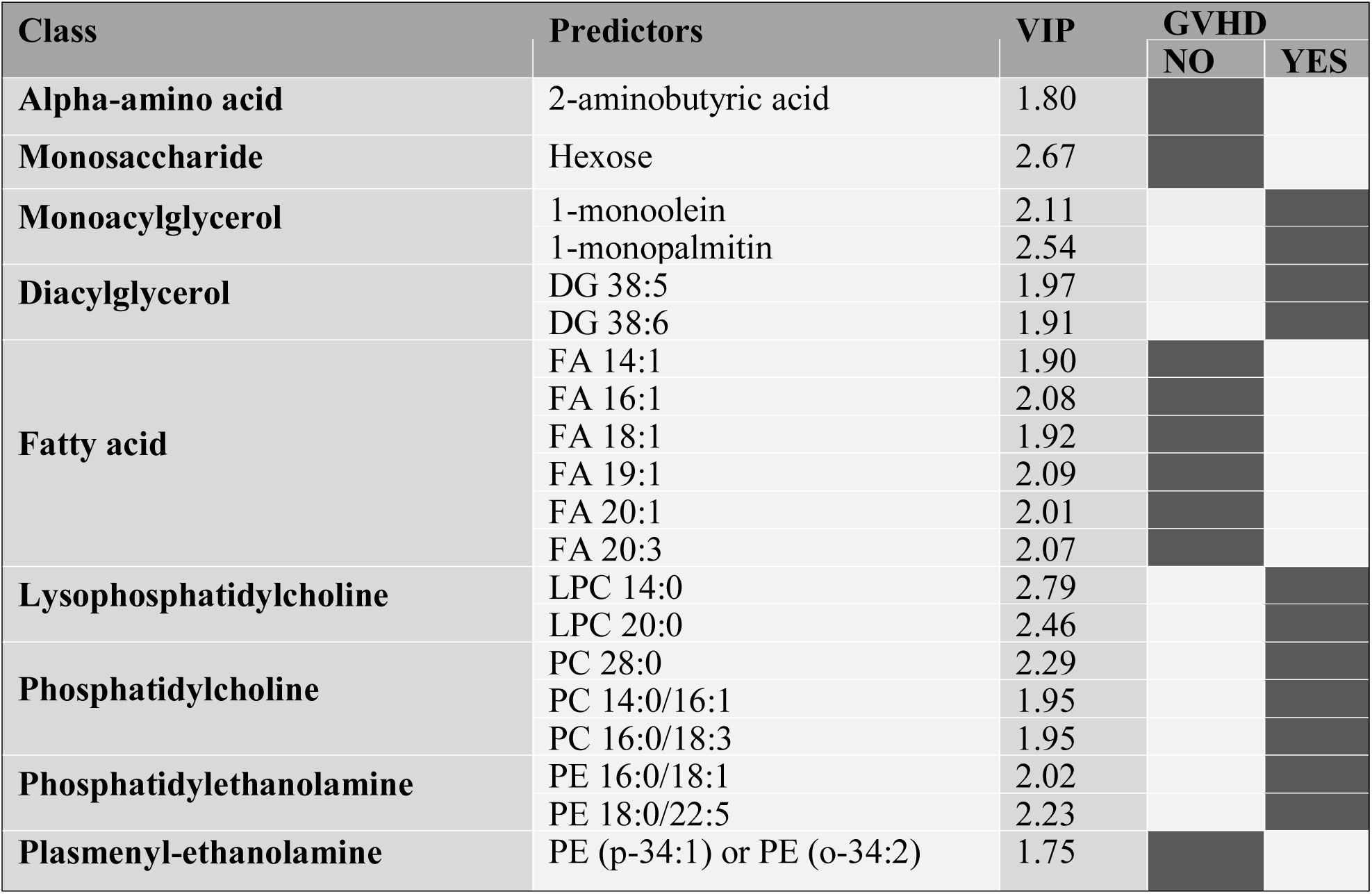
The metabolites and lipids constituting the top 20 VIP’s that predict class separation on the day of transplant between those patients who eventually went on to develop GVHD as opposed to those who did not develop GVHD following BMT. Patients that will develop GVHD had decreased 2-aminobutyric acid, hexose, unsaturated fatty acids, and plasmenyl-ethanolamine PE (p-34:1) or PE (o-34:2), along with elevated monoacylglycerols and diacylglycerols, lysophosphocholines, phosphocholines and phosphoethanolamines. GVHD’ patients (YES) and no-GVHD (NO) are represented by dark grey (high) or light grey (low), respectively.

### The more important variables for class separation suggest metabolic pathway tendencies enabling pre-transplant prediction of onset of GVHD

So far, our study has identified 20 metabolites whose differential presence corelates strongly with the eventual onset of GVHD suggesting an inherent metabolic disturbance that predispose a patient towards GVHD as early as the day of SCT. Further examination of these metabolites indicates that they modulate three related metabolic pathways. Activated phospholipids metabolism appears to be one of the main alterations predicting GVHD pre-transplant. 1-monopalmitin is a monoacylglycerol that had a high VIP score (2.54), and its elevation in GVHD group indicates a metabolic shift in its direction, with phospholipid degradation in cell membranes to produce diacylglycerol, the precursor for MAGs. Alternatively, the no-GVHD state appears to be associated with elevation of monounsaturated and polyunsaturated fatty acids (Figure 3A). Hexose, more commonly called glucose had also a high VIP score (2.67), indicating that elevated glucose uptake is on demand for energy production in the Tricarboxilic Acid Cycle (TCA), as well as increased aerobic glycolysis required for hematopoietic cell proliferation. These process increase the production of NADH used in the electron transport chain, but its upregulation induces excessive reducing power that triggers processes such as fatty acid unsaturation and ROS production (Figure 3B). The levels of oxidative stress originated by excessive ROS production is controled by the glutathione metabolism, where NADH is also used to reduce glutathione for its antioxidant action over ROS. The excessive demand for antioxidative process can deplete glutathione and its precursor cysteine, increasing the demand of 2-aminobutyric acid that can either modulate the glutathione synthesis or be used in ophtalmate synthesis, a tripeptide analog of glutathione, with similar compensative antioxidative actions (Figure 3C).

**Figure 3.**
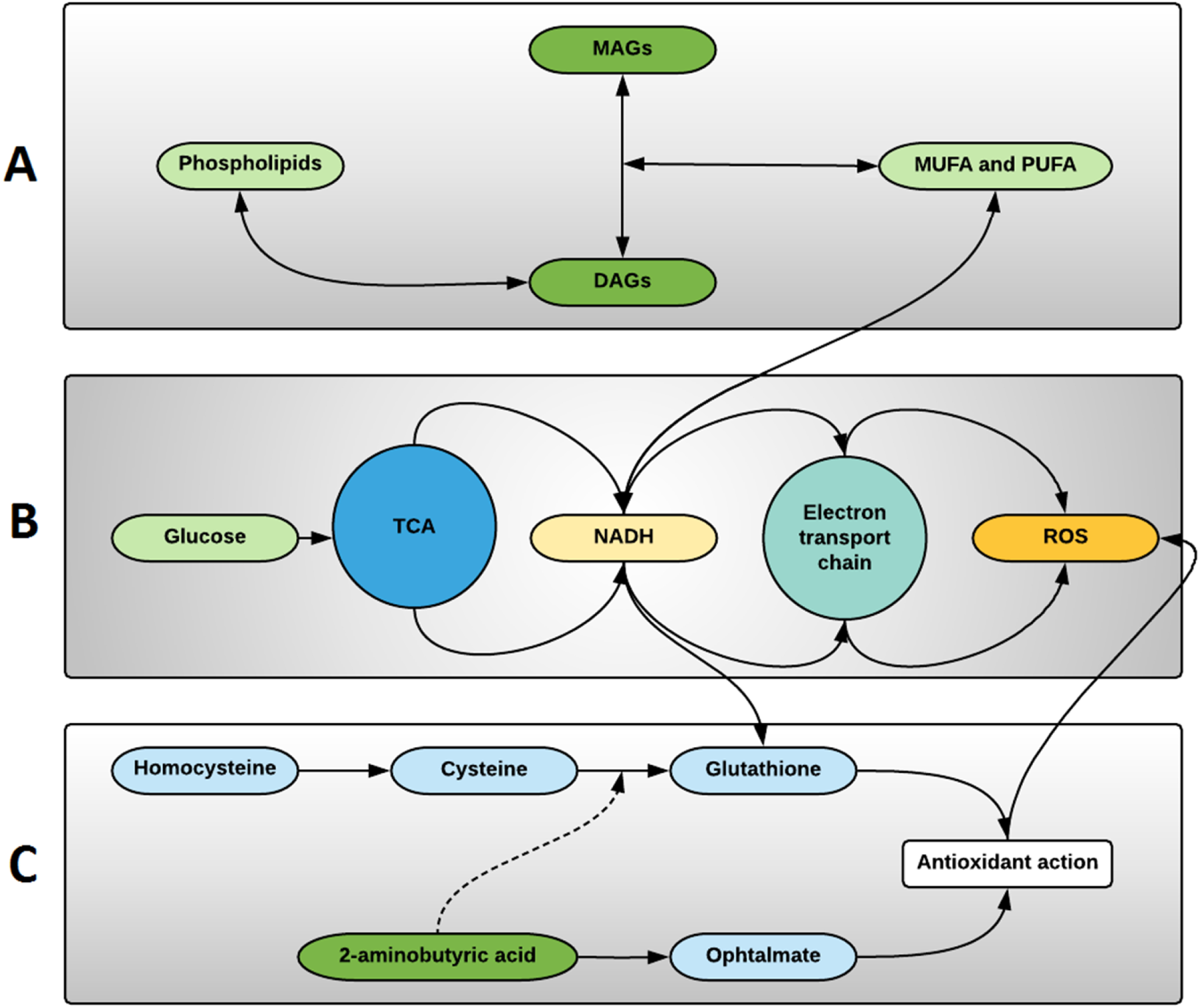
The more important lipids and metabolites predicting GVHD can indicate the main compensatory metabolic pathways modulated in hematopoietic cells pre-transplant. A) Phospholipids metabolism are affected suggesting Phospholipase C activity to produce Diacylglycerols (DAGs), and Diacylglycerol lipase and Monoacylglycerol lipase activity to produce Monoacylglycerols (MAGs)and Free Fatty acids (FFA), respectively. Predominance of monounsaturated (MUFA) and polyunsaturated (PUFA) free fatty acids are result of increased fatty acyl-CoA desaturases activity, increasing the anticancer activity of ω-3 fatty acids. This metabolic pathway is linked to elevated levels of NADH, produced in the Tri-Carboxylic Acid Cycle (TCA), and important on desaturases activity. B) Glucose uptake is increased due to increased energy demands in hematopoietic’ cells, and increased aerobic glycolysis with increased NADH, providing NADH for the electron transport chain and resulting reactive oxygen species (ROS). Oxidative stress compensation is achieved by activation of cysteine-glutathione pathway, and antioxidant action. Exacerbation of these mechanism leads to depletion of cysteine and glutathione, causing increased 2-aminobutyric acid-ophtalmate compensatory pathway activation.

### The more important variables for class separation can be used to build models for GVHD prediction pre-transplant

To evaluate the potential to build models predicting future GVHD the top 20 highest VIP scores were used in an exploratory analysis based in multivariate Receiver Operating Characteristic (ROC) curve method. This method estimated the Area under the Curve of the ROC (AUROC) and bootstrap confidence interval (CI) allowing the selection of best fit model with the desired low data dimensionality. Hence, the analysis showed that plasma metabolites and lipids obtained post-conditioning may be used to build strong predictive models (Figure 4). Models with 2, 3 or 5 variables demonstrated the same level of performance, fitting the selection criteria, with AUCROC ranging from 0.915 to 0.935. With the criteria of finding the lowest number of predictors that can also explain physiologically the metabolic profile of the classes, a model with 5 predictors was chosen for further exploration as potential biomarkers for risk stratification of patients with the potential for development of GVHD following SCT.

**Figure 4.**
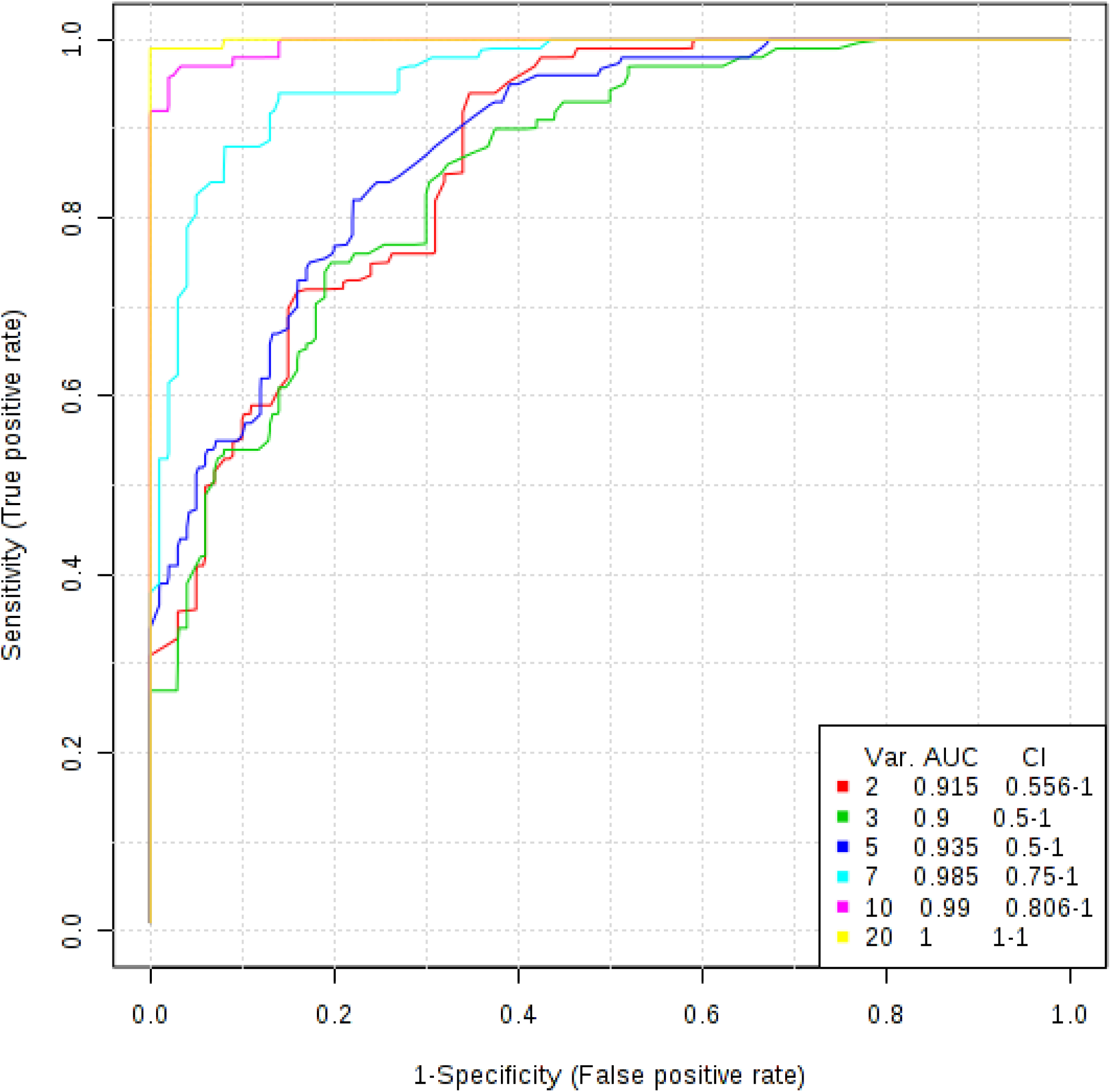
The top 20 variables from PLS-DA method were used to build appropriate classification models with their respective confidence interval. Predictive models pre-transplant demonstrated robust classification even with only two variables.

### Univariate ROC curve analysis finds potential biomarkers of GVHD with plasmatic data pre-transplant

To find biomarkers of GVHD among the top 20 predictors selected in the PLS-DA method, a univariate ROC curve analysis was performed for each of the top 20 variables on day of transplant. The optimal cutoff found for each predictor from the ROC curve was used to estimate the sensitivity and specificity, and to calculate the positive and negative likelihood ratios.

The 5 best biomarkers and their respective estimates are depicted in Table 3, and the comparison plot for each potential biomarker is presented in Figure 5 where presence of outliers is of notice. These data support the previous finding that a model with 5 metabolic biomarkers may provide a robust model for GVHD prediction.

**Table 3.**
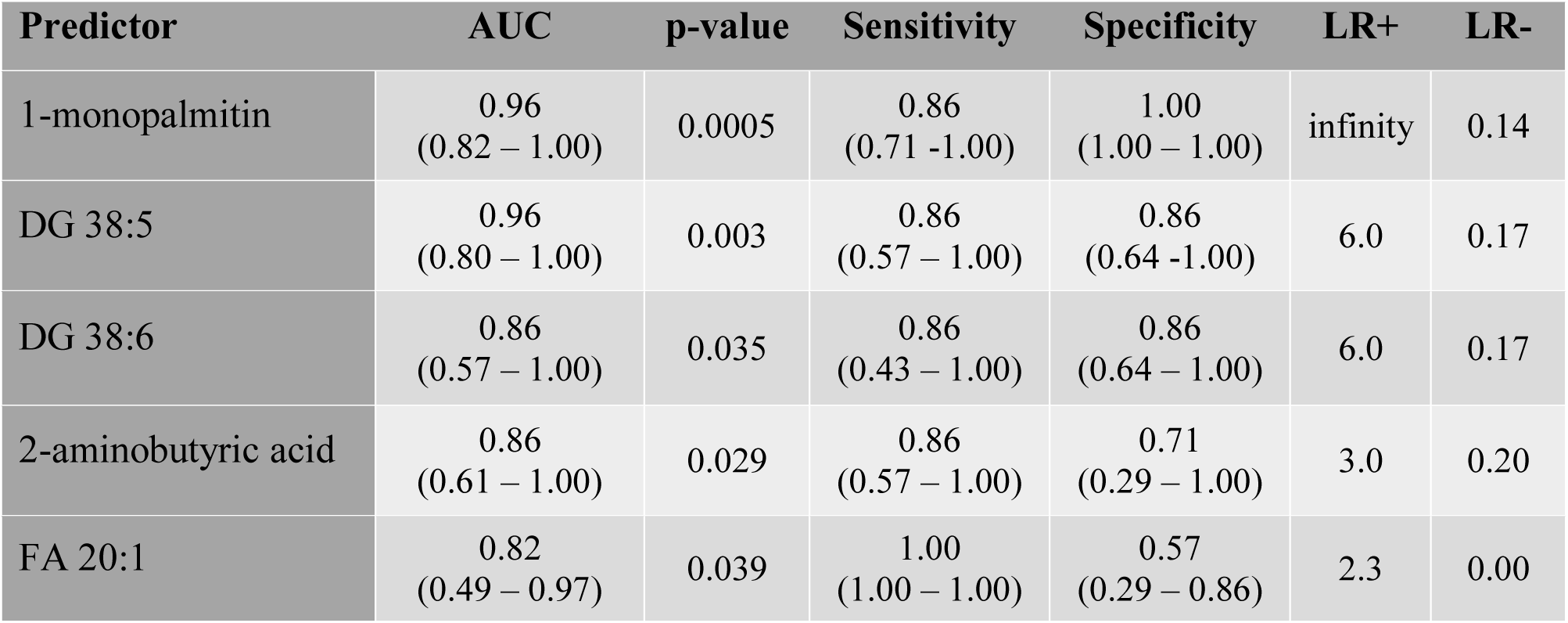
Potential biomarkers performance for a model with top 5 predictors based in accuracy performance (AUC estimate).

| Predictor | AUC | p-value | Sensitivity | Specificity | LR+ | LR- |
| --- | --- | --- | --- | --- | --- | --- |
| 1-monopalmitin | 0.96<br>(0.82 – 1.00) | 0.0005 | 0.86<br>(0.71 -1.00) | 1.00<br>(1.00 – 1.00) | infinity | 0.14 |
| DG 38:5 | 0.96<br>(0.80 – 1.00) | 0.003 | 0.86<br>(0.57 – 1.00) | 0.86<br>(0.64 -1.00) | 6.0 | 0.17 |
| DG 38:6 | 0.86<br>(0.57 – 1.00) | 0.035 | 0.86<br>(0.43 – 1.00) | 0.86<br>(0.64 – 1.00) | 6.0 | 0.17 |
| 2-aminobutyric acid | 0.86<br>(0.61 – 1.00) | 0.029 | 0.86<br>(0.57 – 1.00) | 0.71<br>(0.29 – 1.00) | 3.0 | 0.20 |
| FA 20:1 | 0.82<br>(0.49 – 0.97) | 0.039 | 1.00<br>(1.00 – 1.00) | 0.57<br>(0.29 – 0.86) | 2.3 | 0.00 |

**Figure 5.**
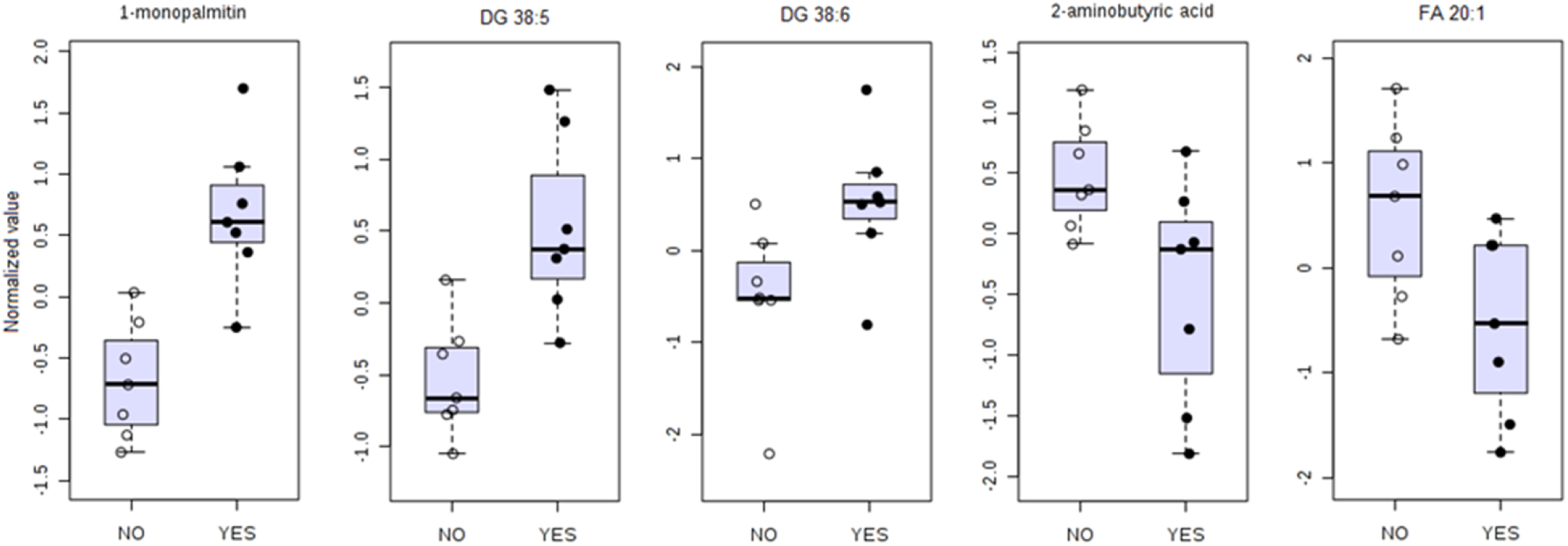
Comparison of 5 biomarkers of GVHD (YES) against no-GVHD (NO) The comparative analysis revealed elevated 1-monopalmitin, DG (38:5) and DG (38:6) and decreased 2-aminobutyric acid and FA (20:1) in patients that will develop GVHD.

All potential biomarkers, shows high sensitivity and specificity, except for FA 20:1 that, while it has maximum sensitivity, has low specificity. The positive likelihood ratio of 1-monopalmitin and DG (38:5) and DG (38:6) shows that a patient with a positive test result for these biomarkers will have very high odds of developing GVHD. 1-monopalmitin, 2-aminobutyric acid, and DG 38:5 showed CI ranging inside an acceptable AUROC values (0.6 – 1.0). Yet, the bootstrap confidence interval for DG 36:6 and FA 20:1 are indicative that these two compounds must be taken with caution despite their high sensitivity estimate.

The evaluation of SVM model performance analyzed by AUROC shows the 5 specific biomarkers model is accurate in predicting the future development of GVHD (AUC=0.995) with patient’s plasma pre-transplant (Figure 6A). The predictive power of class probability of the SVM method was also tested and the confusion matrix represented by the probability plot in Figure 6B shows the method is robust without any misclassification.

**Figure 6.** Final model with 5 specific biomarkers predicting GVHD with cross validation shows strong performance accuracy to predict GVHD (A). The cross-validation for the SVM method for model building and performance evaluation is performed multiple times to test a model with 5 specific biomarkers and it shows that the model can predict GVHD patients without misclassification (B).

## Discussion

The identification of biomarkers that better enable the risk stratification of SCT patients with respect to the potential for development of GVHD has significant clinical utility in the execution of BMT protocols. Such a capability enables the development of new treatments and disease detection and enable evidence based treatments^32,33^. The strategy used in our study to enable this goal investigated the plasma metabolome and lipidome to profile SCT patients. This approach resulted in the generation of complex and multidimensional data that required the use of multivariate statistical approaches to decrease dimensional space, a common approach in metabolomic and lipidomic studies^34,35^. The predictors and potential biomarkers of GVHD found in the plasma of patients pre-transplant indicate that the metabolic profile prior to bone marrow transplant is a major influence in their clinical course following transplantation and can be used to correlate immune and metabolic phenotype. The small size of the cohort used in this study is a limitation that prevents definitive confirmation that the biomarkers identified in this study may be used as predictors to risk stratify patients with respect to the potential for the development of GVHD. Despite this limitation, this study allows an understanding of the metabolic milieu in the study population at the time of transplantation and can be used to direct future studies similar to other studies described in literature^36^. Furthermore, significance following the use of the statistical strategy to overcome the small sample size using Monte Carlo Cross Validation with bootstrap resampling to create confidence intervals further indicate the study results are of significance. Such approaches have been used previously in literature as appropriate tools to deal with the small sample size problem^37^.

The discovery of top 20 lipids and metabolites separating the GVHD from no-GVHD patients identify the main metabolic pathways involved in the phenotype of patients. As an example, 2-aminobutyric acid was identified as a significant biomarker and is a byproduct in the cysteine biosynthesis pathway and relates to glutathione (GSH) metabolism. Soga *et al.*^38^ reported plasma and liver depletion of GSH in acetaminophen poisoning and concomitant increases of ophthalmate. Ophtalmate is a tripeptide (glutamate/2-aminobutyrate/glycine), also known as ophthalmic acid. It is an analog of GSH in which the cysteine group is replaced by L-2-aminobutyrate. It has been proposed that oxidative stress leads to intracellular depletion of GSH, depletion of cysteine, and consequent activation of ophthalmate synthesis. GSH is the predominant low-molecular-weight thiol in animal cells effectively scavenging free radicals. Moreover, its cellular concentration are reduced in response to protein malnutrition, oxidative stress, and many pathological conditions^39,40^. These are all conditions prevalent following myeloablative chemotherapy and radiation therapy. 2-aminobutiric acid also increases intracellular GSH levels by regulation of AMP-activated protein kinase and increase of NADPH and reduction of GSH. Plasma levels of cysteine is also regulated by oxidative stress and that supply of cysteine is directly related to GSH synthesis^41^. We found that cysteine levels are slightly elevated in the GVHD group, although the difference compared with no-GVHD is not statistically significant (p=0.07). This find indicates that pre-transplant the GVHD group profile favors the ophthalmate pathway to compensate for oxidative stress.

The depletion of GSH in cancer cells is one of the targets for chemotherapy, since the increase in basal ROS levels leads to cell death by apoptosis^42^. It has been demonstrated that chemotherapy agents are capable of inactivating glutathione reductase, the enzyme that catalyze the reduction of glutathione disulfide to the sulfhydryl form glutathione, which is a critical molecule in resisting oxidative stress and maintaining the reducing environment of the cell^43,44^. A pitfall of GSH inhibition is that just as cancer cells use GSH activation to modulate levels of ROS, other critical cell populations use this same mechanism to maintain cellular homeostasis^45^. Particularly in the immune system, activated T cells undergoing clonal expansion have increased energy demands that increase production of ROS by the mitochondrial electron transport chain^46^. GSH is not necessary for cell activation, but activated T cells regulate their oxidative stress by using GSH, a key component for metabolic reprograming for cell differentiation^47^. TCR ligation and binding with costimulatory molecules induces metabolic remodeling of the naive T cell to anabolic growth and biomass accumulation, and increases aerobic glycolysis, even though sufficient oxygen is present to support glucose catabolism via the tricarboxylic acid cycle^30^. This phenomenon is termed the Warburg effect and also happens in proliferating cancer cells. T cells do not need aerobic glycolysis for activation, but during continuous proliferation, along with oxidative phosphorylation, the process is triggered and is vital for T cells effector functions, as cytokine production^48^. Therefore, high energy demand in the proliferative phase leads to increased glucose uptake and accelerated glucose metabolism, enhance aerobic glycolysis besides oxidative phosphorylation. Patients that will eventually develop GVHD appear to demonstrate this metabolic profile on the day of stem cell, and donor T cell infusion.

Another aspect of the metabolic profile of GVHD-prone patients is that lipid metabolism is significantly altered, placing lipids as the main biomarkers to predict GVHD. Lipid modulation is expected due to metabolic demands of compromised hematopoietic tissue, immunologic response and underlined inflammatory profile of the patients related to the conditioning regimen. Of the important plasma biomarkers of GVHD, free fatty acids found in our study are monounsaturated (MUFA) and polyunsaturated fatty acids (PUFA). The effects of MUFA and PUFA from the omega-3 family in decreasing inflammation have been extensively studied in effort to introduce intake of dietary lipids toward treatment of several diseases^49^. In addition to modifying the profile of eicosanoids involved in inflammatory processes, this lipid family affects the production of many inflammatory proteins including cytokines as well as, pro-inflammatory tumor necrosis factor (TNF-α), interleukin IL-1 β and IL-6. These also have the potential to decrease production of these cytokines in response to LPS and increase the concentration of the anti-inflammatory cytokine IL-10^50^. MUFA and PUFA are produced by the desaturation process involving three broad specificity fatty acyl-CoA desaturases that introduce unsaturation at C5, C6 or C9 position. Stearoyl-CoA desaturase (SCD) is the rate-limiting enzyme catalyzing the synthesis of monounsaturated fatty acids (MUFAs), primarily oleic acid (C18:1) and palmitoleic acid (C16:1). The ratio of saturated to monounsaturated fatty acids in membrane phospholipids is critical to normal cellular function and alterations in this ratio have been correlated with diabetes, obesity, cardiovascular disease, and cancer, and the oxidative stress characteristic of these pathologies^51^. The electrons transferred from the oxidized fatty acids during desaturation are transferred from the desaturases to cytochrome b5 and then NADH-cytochrome b5 reductase. These electrons are uncoupled from mitochondrial oxidative-phosphorylation and, therefore, do not yield ATP. Moreover, electrons can leak from the respiratory chain during oxidative metabolism, and partially reduce molecular oxygen into ROS, contributing to oxidative stress^52^. This oxidative stress at the critical time of donor T cell infusion is likely related to GVHD.

In a study analyzing plasma phospholipids changes in patients with acute leukemia, it was demonstrated that all phospholipids’ concentrations found in patients at the time of diagnosis were significantly lower than in reference group^53^. The phospholipid down-regulation is an important scenario to understand the effect of these lipids in predicting GVHD. Endogenous lipids are important lipids not only in regulation of inflammation, but also in expressing antitumor functions in several types of cancer^54–56^. Free fatty acids may be esterified in cell membrane phospholipids undergoing hydrolysis by phospholipases to generate bioactive lipid mediators, including lysophosphocholine (LPC), diacylglycerol (DAG), monoacylglycerol (MAG), and unsaturated fatty acids (MUFA and PUFA). LPC is known to exert immune-regulatory activity, by increases in the numbers of T cells, monocytes and neutrophils, and also inducing protein kinase C activation in bone marrow derived mastcells^57^. In an in vitro study in vascular smooth muscle cells, treatment with LPC caused a 2-fold increase in intracellular ROS that was blocked by the NADH/NADPH oxidase inhibitor. This indicates that ROS generated by NADH/NADPH oxidase contribute to LPC-induced activation of extracellular signal-regulated kinase 1/2 (ERK1/2), and stimulates the growth and migration of vascular smooth muscle cells^58^. DAGs are not only a precursor for free fatty acids but also an important signaling molecule in cells. Protein kinase C (PKC) is the major cellular target of DAG and its binding leads to the PKC plasma membrane translocation and activation. The protein kinase D (PKD) is a substrate of PKC and after activation moves among different subcellular compartments, being responsible for several cell responses as proliferation, differentiation, apoptosis, and immune response through TCR signalling^59^. Under oxidative stress, ROS induces activation of PKD, which activates the transcription factor NF-kB to protect the cell from oxidative-stress-induced cell death^60^. 1-monopalmitin, also called 1-palmitoyl-sn-glycerol is a monoacylglyceride formed via release of a fatty acid from diacylglycerol by diacylglycerol lipase. MAGs have also been indicated as signaling bioactive molecules. The hydrolysis of (MAG) to FFA and glycerol, is conducted by the MAG lipase (MAGL) in different tissues, although ABHD6, a MAG hydrolase, has been implicated in the pathogenesis of metabolic syndrome^61^, inflammation^62^, and in cancer^63^. Deletion or inhibition of ABHD6 activity has been shown to be beneficial in certain cancers^64^. The importance of 1-monoacylglycerols (1-MAGs) with a saturated fatty acid group, is demonstrated by its accumulation upon ABHD6 suppression, and direct binding to the ligand binding domain of PPARα and PPARγ, and activating these transcription factors^65^. MAGs metabolism is also related to the effects of endocannabinoids in the immune system. The endocannabinoid receptor CB2 is identified as a peripheral receptor preferably present in B cells, T cells, macrophages, monocytes, natural killers and polymorphonuclear cells^66^. The enzyme MAGL that hydrolyzes of MAGs also hydrolyzes 2-arachidonoylglycerol (2-AG), an endogenous endocannabinoid acting through CB2 receptors in the immune system with immune suppression effects^67^.

## Conclusion

Our study demonstrates that the pre-transplant lipidome and the metabolome of SCT recipients has significant potential towards their risk stratification with respect to the development of the GVHD indicating potential use as biomarkers for this purpose. The identified potential biomarkers indicate a pro-inflammatory metabolic profile in patients that will eventually develop GVHD. The role of GSH and its association with 2-aminobutyric acid, coupled with signs of altered glucose metabolism, support the hypothesis that in patients that will develop GVHD, GSH levels are excessively depleted due to elevated oxidative stress related to glucose metabolism, in response to chemotherapy treatment. In this scenario both groups have elevated 2-aminobutyric acid as a compensatory mechanism, although in patients susceptible to develop GVHD it is not produced in a large enough amount to compensate for the increased antioxidant demand. Furthermore, the decreased levels of plasma hexose indicates excessive glucose uptake for glycolysis and oxidative phosphorylation, and consequent ROS production. The protective effect of MUFA and PUFA as anti-inflammatory agents is decreased in patients that will develop GVHD and elevation of phospholipids and DAG and MAG indicates an increased traffic of inflammatory lipids. This pro-inflammatory profile in patients with risk of GVHD is associated with immune suppression and characterize an unfavorable environment coupled with the overwhelming physiologic impact of undergoing the donor graft and predisposes the host to GVHD.

## Acknowledgement

Research reported in this publication was supported by research grants from National Institutes of Health under grant numbers HD087198 (to DSW). The content is solely the responsibility of the authors and does not necessarily represent the official views of the National Institutes of Health. This work also received support via a Young Investigator Award from SCIEX for clinical lipidomic research (DSW).

## Statement of Competing Financial Interests

All authors of this manuscript declare that they have no competing financial interests.

